# A cataract-causing Y204X mutation of CRYßB1 promotes C-terminal degradation and higher-order oligomerization

**DOI:** 10.1101/2023.02.24.529791

**Authors:** Xuping Jing, Xiaoyun Lu, Mingwei Zhu, Lingyu Shi, Ping Wei, Bu-Yu Zhang, Yi Xu, Dao-Man Xiang, Ya-Ping Tang, Peng Gong

**Author notes:** Address correspondence to: Peng Gong, Ya-Ping Tang, Dao-Man Xiang.

## Abstract

Crystallin (Cry) proteins are a class of main structural proteins of vertebrate eye lens, and their solubility and stability directly determine transparency and refractive power of the lens. Mutation in genes that encode for these Cry proteins is the common cause for congenital cataract. Despite extensive studies, the pathogenic and molecular mechanisms remain unclear. In this study, we identified a novel mutation in *CRY*_Β_*B1* from a congenital cataract family, and demonstrated that this mutation led to an earlier termination of protein translation, resulting in a 49-residue truncation at the CRYβB1 C-terminus. This mutant is susceptible to proteolysis and allows us to determine a 1.2- Å resolution crystal structure of CRYβB1 without the entire C-terminal domain. In this crystal lattice, two N-terminal domain monomers form a dimer that structurally resembles a wild-type (WT) monomer, but with different surface characteristics. Biochemical analyses suggest that this mutant is significantly more liable to aggregate and degrade, when compared to WT CRYβB1. All our results provide an insight into the mechanism regarding how a mutant Cry contributes to the development of congenital cataract possibly through alteration of inter-protein interactions that result in the opacity of eye lens.

## Introduction

Cataract, the opacity or light scattering of eye lens caused usually by the presence of abnormal crystallin (Cry) protein aggregates, is the main cause of human blindness worldwide. It can be broadly divided into congenital and acquired cataract, with the congenital cataract being one of the major causes of childhood blindness (1-3). Crystallin proteins are the major structural components of the vertebrate eye lens, and their solubility and stability are important for maintaining transparency and refractive power of the eye lens (1). They can be grouped into two families, the α-crystallins superfamily and β/γ-crystallins superfamily (4). Human β- and γ-crystallins share highly homologous amino acid sequences and common structural feature consisting of an N-terminal domain (NTD) and a C-terminal domain (CTD) connected by a short linker (5). These two domains are believed to evolve from a common single-domain protein ancestor by a series of gene duplication and fusion events (6,7). Each domain consists of two similar highly stable Greek key motifs of about 40 amino acids, folding into a wedge-shaped β-sheet sandwich with a pseudo 2-fold symmetry (1). The main sequence difference between oligomeric β-crystallins and monomeric γ-crystallins is that only the former has long N-terminal extensions. The terminal extensions of β-crystallins are thought to be involved in the formation and stabilization of higher-order homologous or heterogeneous oligomer (8-10). The NTD and CTD of β/γ-crystallins arrange as an inter-molecularly domain-swapped dimer or intra-molecularly face-to-face dimer according to available structures in the protein data bank (1,5,6,11-14). The structural integrity of the Greek key motifs in β/γ-crystallins is vital for proper folding of protein, and mutations disrupting even one of the four Greek key motifs would result in protein self-aggregation and precipitation, consistent with the phenotype of nuclear cataract (4,15-17). Correspondingly, mutation of crystallin genes is a dominant pathogenic factor of the congenital cataract characterized by lens opacity (18).

β-crystallins can be further classified into basic and acidic groups. Basic crystallins (βB1, βB2, and βB3) contain both N- and C-terminal extensions, while acidic crystallins (βA1, βA2, βA3 and βA4) possess only N-terminal extension (5). CRYβB1 is a primary member of β-crystallins and comprises 9% of the total soluble crystallin proteins in the human lens (17,18). Clinically identified mutations (p.Q227X, p.Q223X, p.G220X, p.X253R, p.S228P, p.R233H, p.S93R, and p.S129R; each X denotes a translation termination codon and results in a C-terminal truncation of the protein) in CRYβB1 were found to be associated with the autosomal dominant cataract (4,16,17,19-23). The C-terminal truncations lead to partial or complete deletion of the C-terminal extension, or even affect the integrity of the CTD. For documented autosomal dominant congenital cataract-related CRYβB1 truncation-type mutations (p.Q227X, p.Q223X and p.G220X), it has been proposed that the mutations disrupt the hydrophobic core of the CTD, leading the mutants to be insoluble and easier to aggregate and eventually causing cataract (4,16,20). However, the molecular mechanism underlying the pathogenesis of congenital cataract remains unclear. The three-dimensional structure and functional exploration of novel congenital cataract-related crystallin mutants could provide important references in elucidating the pathogenic mechanism, and therefore beneficial for translational research efforts on curing congenital cataract.

In this study, we identified a previously unreported truncation-type mutation (nucleotide: c.612 C>A; amino acid: p.Y204X) in exon 6 of *CRY*_Β_*B1*, a cataract-causative gene in a family with autosomal dominant congenital cataract (ADCC). To explore the molecular mechanism of the pathogenic role, we solved a 1.20-Å resolution crystal structure of the mouse CRYβB1 truncated mutant Y202X (M_Y202X, the corresponding mutation of human p.Y204X in mouse, which also causes cataract in mouse; data not shownequivalent to the human CRYβB1 Y204X). The structure only contains the CRYβB1 NTD, but forms WT CRYβB1 monomer-mimicking dimer in the crystal lattice. Furthermore, the mutant protein shows different structural and biochemical characteristics when compared to those in WT CRYβB1, with respect to oligomerization and proteolysis-related degradation. This work thus provides key information for further understanding of the molecular mechanism of this and similar pathogenic mutation.

## Results

### Identification of a novel *CRY*_Β_*B1* gene mutation associated with cataract in a Chinese family

In a three-generation Chinese family, four members suffered from nuclear cataract, with an obvious hereditary tendency (Figure 1, A-B). To identify the genetic factors responsible for cataract in this family, whole-exome sequencing was performed with gDNAs from patients and their relatives. A heterozygous *CRY*_Β_*B1* c.612 C>A mutation (NM_001887.4), which was not reported previously, was detected in all four patients (III:4, III:5, III:6, and III:7) and two unaffected members (II:3 and II:5) (Figure 1C). This mutation transited the Y204 codon (TAC) to a termination codon (TAA), resulting in an earlier termination of translation, and thereby produced a truncation-type CRYβB1 Y204X mutant without C-terminal 49 amino acid residues (residues 204-252), in which the Greek key motif IV and C-terminal extension was included. As II:3 and II:5 carried the mutation but did not show cataract, we considered it a causative genetic factor with incomplete dominance. Next, to further confirm the pathogenicity of the mutation and to illustrate the function of the C-terminal extension, we set out to explore the structure and biochemical properties of the truncated CRYβB1 protein.

**Figure 1.**
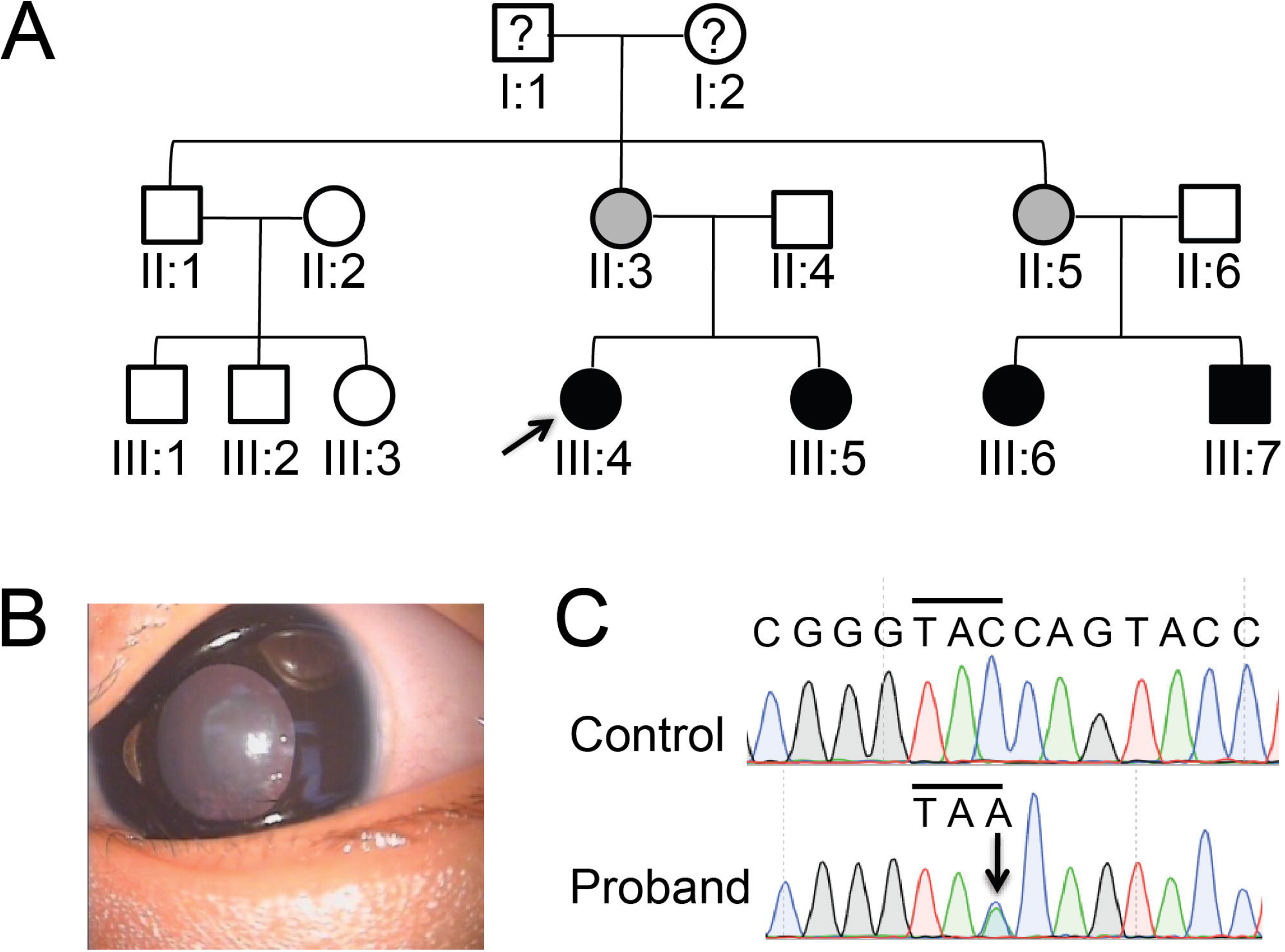
The Mutation analysis of *CRY*_Β_*B1* in a Chinese family with congenital nuclear cataract. A) Pedigree of the family. Squares and circles indicate men and women, respectively. Solid symbols denote affected status and the grey symbols denote the normal people with the mutated gene. The proband is indicated with an arrow. The question mark denotes the person without sequencing. B) Slit-lamp view of the lens of the proband shows the nuclear cataract. C) The DNA sequence chromatogram shows a C612A heterozygous mutation in *CRY*_Β_*B1* indicated by an arrow.

### Crystallization and structure determination of the mouse CRYβB1 Y202X mutants

The N-terminal hexahistidine-tagged human WT CRYβB1 (H_WT), human Y204X mutant (H_Y204X), mouse WT CRYβB1 (M_WT), and mouse CRYβB1 M_Y202X mutant were purified and then subjected to crystallization trials (Figure 2, A, B, and E). Block shape crystals of the M_Y202X were obtained in initial crystallization screening after one week while crystallization of the other three proteins was unsuccessful. The best M_Y202X single crystals were at least 0.1 mm in each dimension and allowed us to obtain an X-ray diffraction data set at 1.20-Å resolution. As the sequence identity between the mouse and human CRYβB1 proteins is 80% with 95% residue coverage, we chose a human CRYβB1 structure (1) as the template to generate molecular replacement (MR) search models (Figure 2C). Successful MR was achieved with a modified NTD model that contains 88 residues (human CRYβB1 residues 54-141). Silver staining of the crystal samples indicated that the molecular weight of the crystallized M_Y202X protein is in the range of 10-15 kDa, suggesting degradation of the protein (Figure 2, E-F). R_work_/R_free_ values after the initial round of structure refinement were 0.204/0.216 (Table 1). After reiterative model building and refinement trials, the final structure model had R_work_/R_free_ values of 0.185/0.194 with 91 residues resolved (Table 1). The crystallographic asymmetric unit comprises a dimer with nearly identical conformation. The root mean square deviation (RMSD) value for all α-carbon atoms is 0.6 Å with 96% coverage (chain A as the reference), and the arrangement of the two NTD monomers mimics one WT protein monomer. The RMSD value for all superimposable α-carbon atoms is 1.1 Å with 93% coverage (mutant structure as the reference, Figure 2D). Despite the observed dimeric form in the crystal lattice, the M_Y202X primarily exists as high molecular weight species in gel-filtration chromatography (Figure 2B).

**Figure 2.**
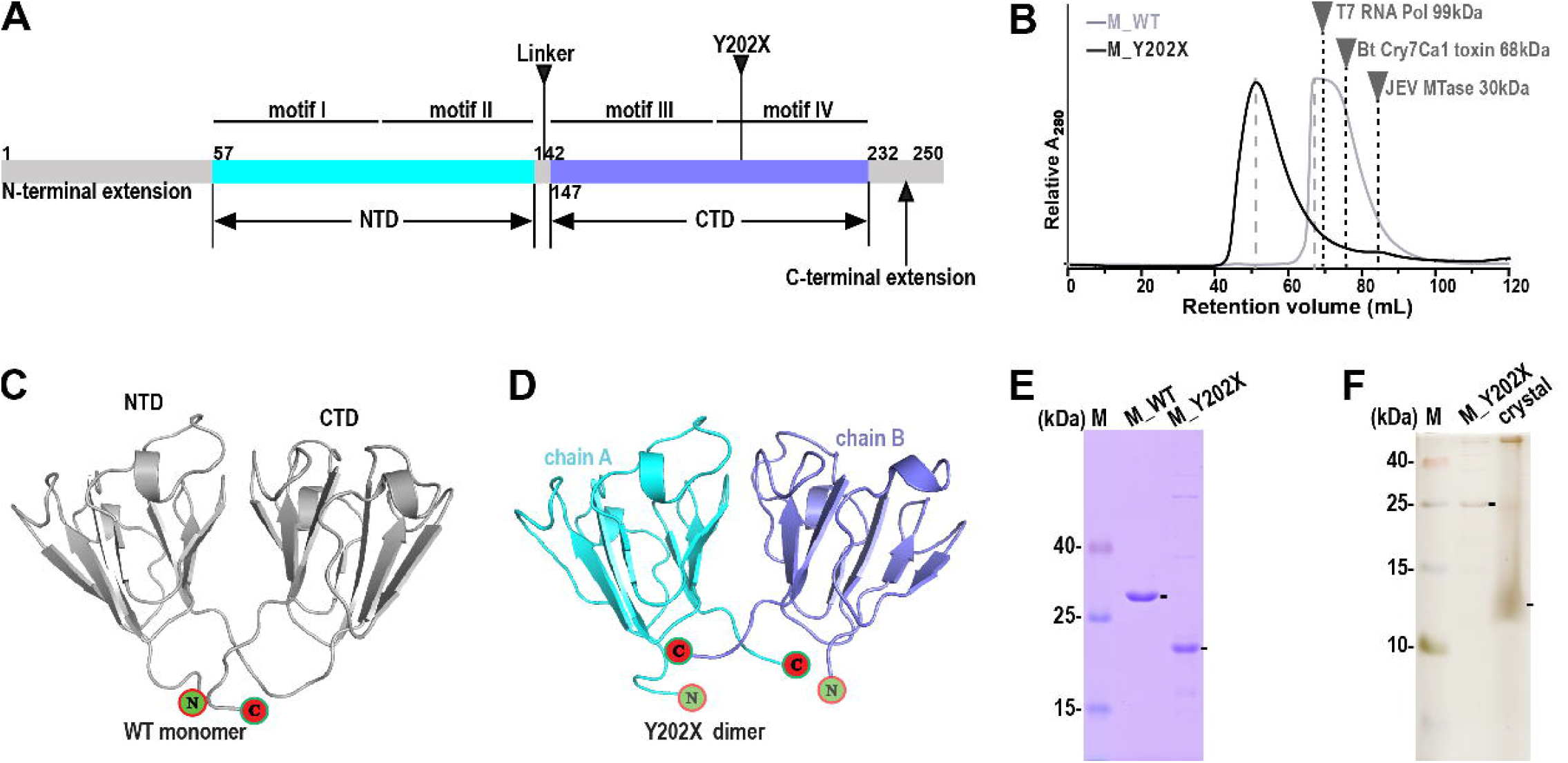
Overall structure of the CRYβB1 mutant. A) A color-coded bar defining structural elements of the mouse CRYβB1 WT protein. The N-terminal extension (grey), NTD (cyan), the linker (grey), the CTD (slate) and the C-terminal extension (grey) are individually color-coded. The four motifs and the mutation site Y202X are indicated according to the residue region or site. B) M_Y202X exhibited a high molecular weight polymeric state in a superdex 200 gel filtration column, while the M_WT exists probably as a tetramer according to the retention volume. Empirical retention volumes and the molecular weights of three other proteins are indicated for comparison. JEV MTase: Japanese encephalitis virus methyltransferase. T7 RNA Pol: T7 RNA polymerase. Bt: *Bacillus thuringiensis*. C) The monomer structure of the human CRYβB1 WT protein (PDB: 1OKI). The whole structure is colored in grey70. D) The M_Y202X dimer structure. Chain A is colored in cyan, and chain B in slate. The N- and C-termini are shown by circles filled with green and red backgrounds, respectively. E) The purified M_WT and M_Y202X proteins with relative higher purify. F) The M_Y202X crystal analysis in Tris-Tricine-SDS-PAGE. The M_WT, M_Y202X and crystal bands are indicated by short lines.

**Table 1.**
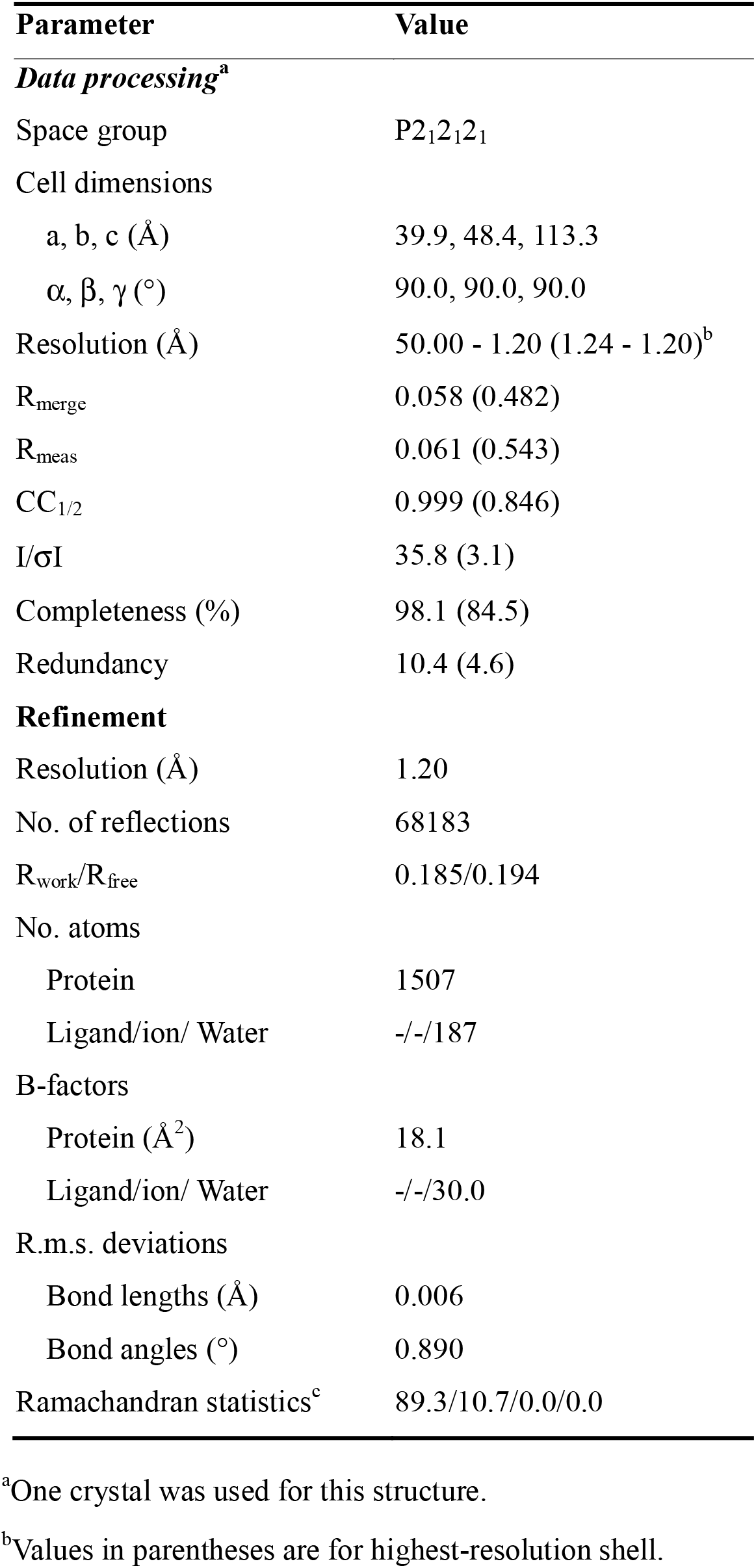
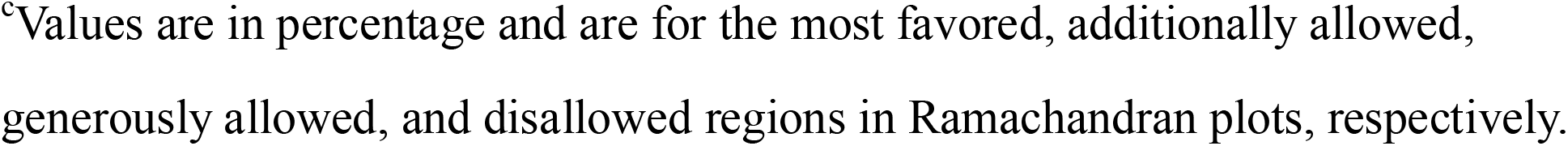
X-ray diffraction data processing and structure refinement parameters.

| Parameter | Value |
| --- | --- |
| *Data processing*<sup>a</sup> |  |
| Space group | P2<sub>1</sub>2<sub>1</sub>2<sub>1</sub> |
| Cell dimensions |  |
| a, b, c (Å) | 39.9, 48.4, 113.3 |
| α, β, γ (°) | 90.0, 90.0, 90.0 |
| Resolution (Å) | 50.00 - 1.20 (1.24 - 1.20)<sup>b</sup> |
| R<sub>merge</sub> | 0.058 (0.482) |
| R<sub>meas</sub> | 0.061 (0.543) |
| CC<sub>1/2</sub> | 0.999 (0.846) |
| I/σI | 35.8 (3.1) |
| Completeness (%) | 98.1 (84.5) |
| Redundancy | 10.4 (4.6) |
| **Refinement** |  |
| Resolution (Å) | 1.20 |
| No. of reflections | 68183 |
| R<sub>work</sub>/R<sub>free</sub> | 0.185/0.194 |
| No. atoms |  |
| Protein | 1507 |
| Ligand/ion/ Water | -/-/187 |
| B-factors |  |
| Protein (Å<sup>2</sup>) | 18.1 |
| Ligand/ion/ Water | -/-/30.0 |
| R.m.s. deviations |  |
| Bond lengths (Å) | 0.006 |
| Bond angles (°) | 0.890 |
| Ramachandran statistics<sup>c</sup> | 89.3/10.7/0.0/0.0 |

<sup>a</sup>One crystal was used for this structure.540 <sup>b</sup>Values in parentheses are for highest-resolution shell.
<sup>c</sup>Values are in percentage and are for the most favored, additionally allowed, generously allowed, and disallowed regions in Ramachandran plots, respectively.

### Overall structure of the mouse CRYβB1 Y202X mutant and a comparison with WT protein structure

The two M_Y202X NTD monomers pair in a symmetrical manner very similar to that observed in the CRYβB1 H_WT crystallin, where its NTD and CTD (sharing ∼33% sequence identity in mouse) are related by a pseudo 2-fold symmetry (Figure 2, A, C, and D). The M_Y202X NTD structures are almost identical to the NTD of the WT protein (RMSD values for 87-88 superimposable α-carbon atoms: 0.5-0.6 Å, chain A of the mutant structure as the reference) (Figure 2C and 2D). The inter-molecular interaction interface between the M_Y202X NTD monomers is highly similar to the intra-molecular NTD-CTD interaction interface of the H_WT. Both interfaces are featured by hydrogen bonding, charged, and hydrophobic interactions (Figure 3). Two hallmark pairs of arginine and glutamic acid form salt bridges in M_Y202X NTD dimer and the H_WT monomer, respectively, while a notable difference is that a pair of R130 residues (one from each NTD monomer) interacts with each other at the edge of the interface in the mutant structure (Figure 3).

**Figure 3.**
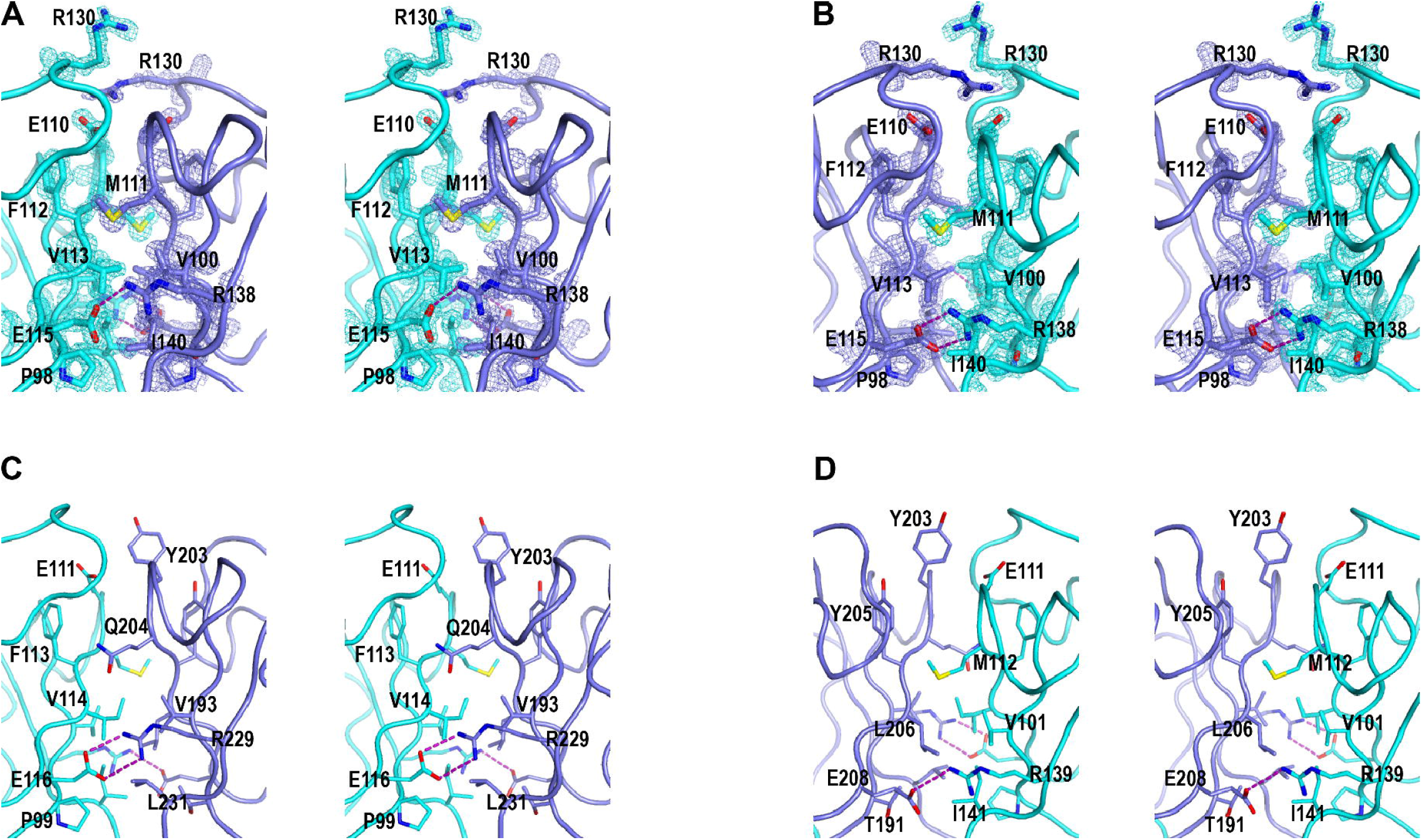
The interface comparison of the CRYβB1 M_Y202X dimer and M_WT monomer. A) The inter-molecular interface of the CRYβB1 M_Y202X dimer. B) The view turn 180° in y-axis from panel A. C) The intra-molecular interface of the CRYβB1 M_WT monomer. D) The view turn 180° in y-axis from panel C. Stereo-pair images of the composite SA omit electron density map (contoured at 1.5 σ, see experimental procedures) overlaid onto structural models around the inter-molecular interface of M_Y202X dimer. The M_Y202X dimer is colored as figure 1D. The NTD and CTD of WT protein shown in thinner representations are also colored in cyan and slate, respectively. The residues located in the exterior of the stereo structures are shown as sticks and labeled. The hydrogen bonds formed by arginine and glutamic acids are shown as purple dashed lines (distance range: 2.5-3.5 Å).

### Surface characteristics of the mouse CRYβB1 mutant Y202X

To compare the surface characteristics between the M_Y202X NTD dimer (M1 and M2) and the WT monomer (NTD and CTD), a M_WT protein model was obtained using the PhyRe2 server (24). Though the M_Y202X NTD dimer can well mimic the WT protein monomer in shape, it shows different electrostatic features in the solvent exposed surface of the second NTD monomer (M2, corresponding to the WT CTD in the comparison), with M2 displaying overall opposite electrostatic potentials in the top, side and bottom regions (Figure 4). These differences could help understand the low solubility and aggregation of this mutant.

**Figure 4.**
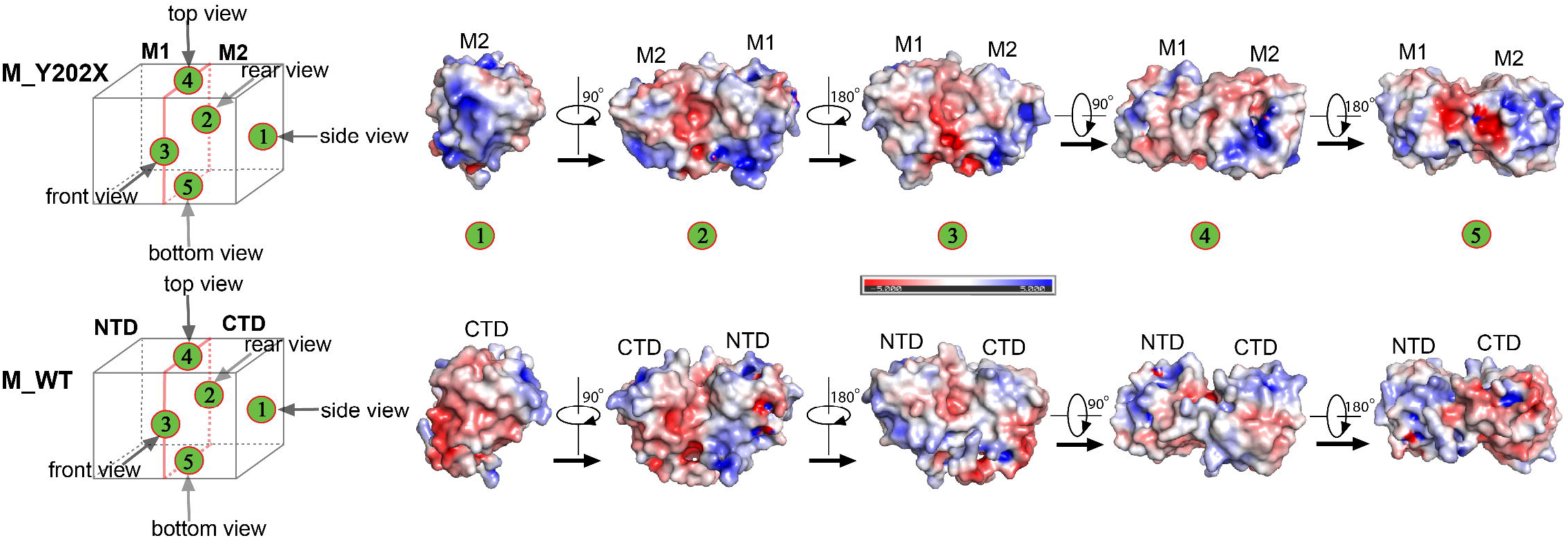
A surface characteristic comparison of the CRYβB1 mutant M_Y202X and M_WT protein. A diagram showing different views of M_Y202X or M_WT is drawn to distinguish the different surfaces more easily. The two domains (M1 and M2, NTD and CTD) are divided by red solid and dotted lines. The 1, 2, 3, 4 and 5 in circles filled with green and red backgrounds denote the side, rear, front, top and bottom views, respectively. The top panel is the M_Y202X and the bottom panel is the M_WT.

### The mutants are more liable to aggregate in solution than WT proteins

In order to further explore the effects brought by the truncation-type mutation regarding solution properties, we analyzed the WT and M_Y202X and the H_Y204 mutants by several biochemical methods. The gel filtration chromatography data showed that the both mutants are high molecular weight polymers, while both WT protein are low molecular weight oligomers according to their retention volumes in a superdex 200 gel filtration column (Figure 2B). The dynamic light scattering (DLS) data indicated that the hydrodynamic sizes of both mutants in phosphate buffer saline (PBS) are larger than those of the WT proteins at 25°C (Figure 5A). Moreover, the negative stain electron microscopy (EM) data also showed that the sizes of the mutant particles are inhomogeneous and much larger than the WT proteins (Figure 5B). The propensity of the mutants to form high molecular weight (HMW) aggregates is compatible with the clinical observation that truncation-type mutations lead to the human congenital cataract.

**Figure 5.**
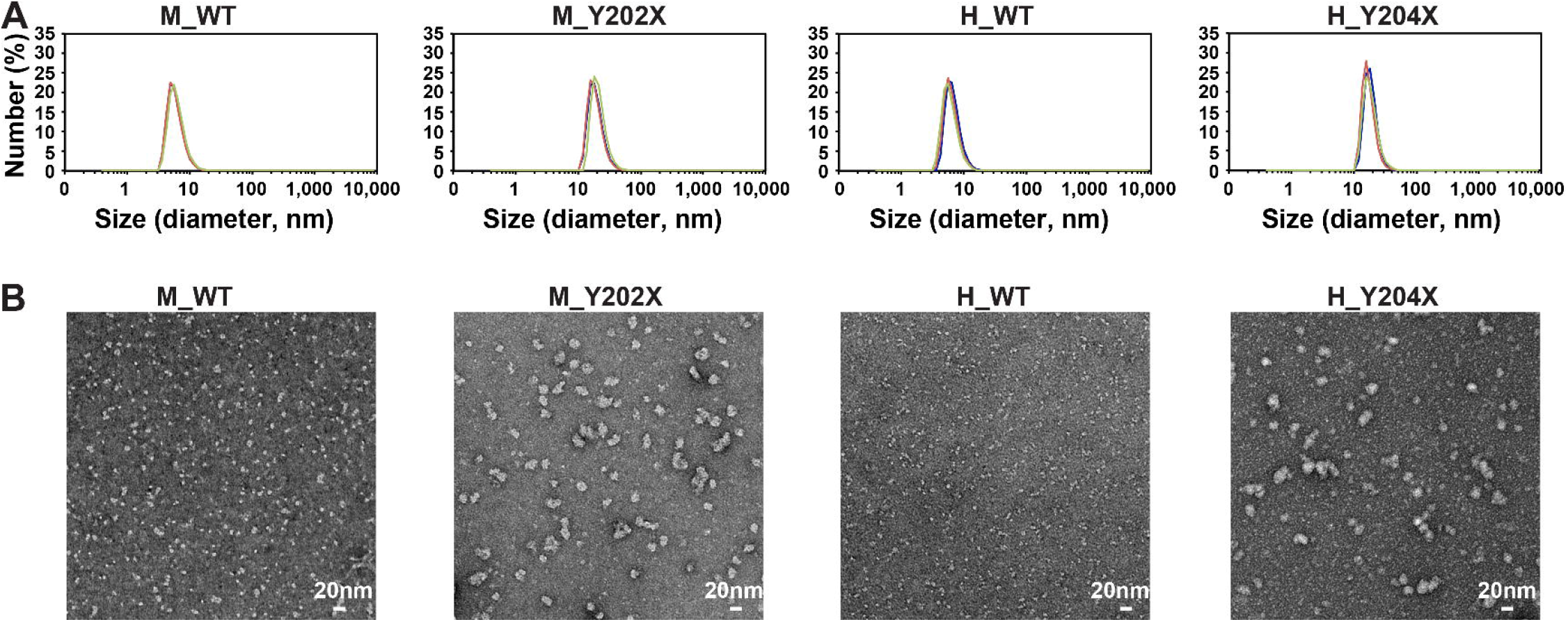
The aggregation formation analysis of CRYβB1 WT and mutated proteins. A) Distribution of protein hydrodynamic diameters in PBS measured by DLS. The peaks of the M_Y202X and H_Y204X are larger than the WT proteins. B) Representative negative-staining electron microscopy micrographs of the CRYβB1 proteins. Scale bar, 20 nm.

### The mutants are more sensitive to the trypsin than WT proteins

The M_Y202X and H_Y204X mutants are susceptible to degradation in the process of protein expression and purification. In order to assess the degradation behaviors of the mutants in solution, trypsin proteolysis analyses were performed. Both mutants were degraded to higher degrees than the WT proteins under the same proteolysis condition (Figure 6, lanes 4-15 for M_WT, lanes 19-30 for M_Y202X, lanes 34-45 for H_WT, lanes 49-60 for H_Y204X). These data suggest that both mutations lead to enhanced protein sensitivity to trypsin proteolysis in solution.

**Figure 6.**
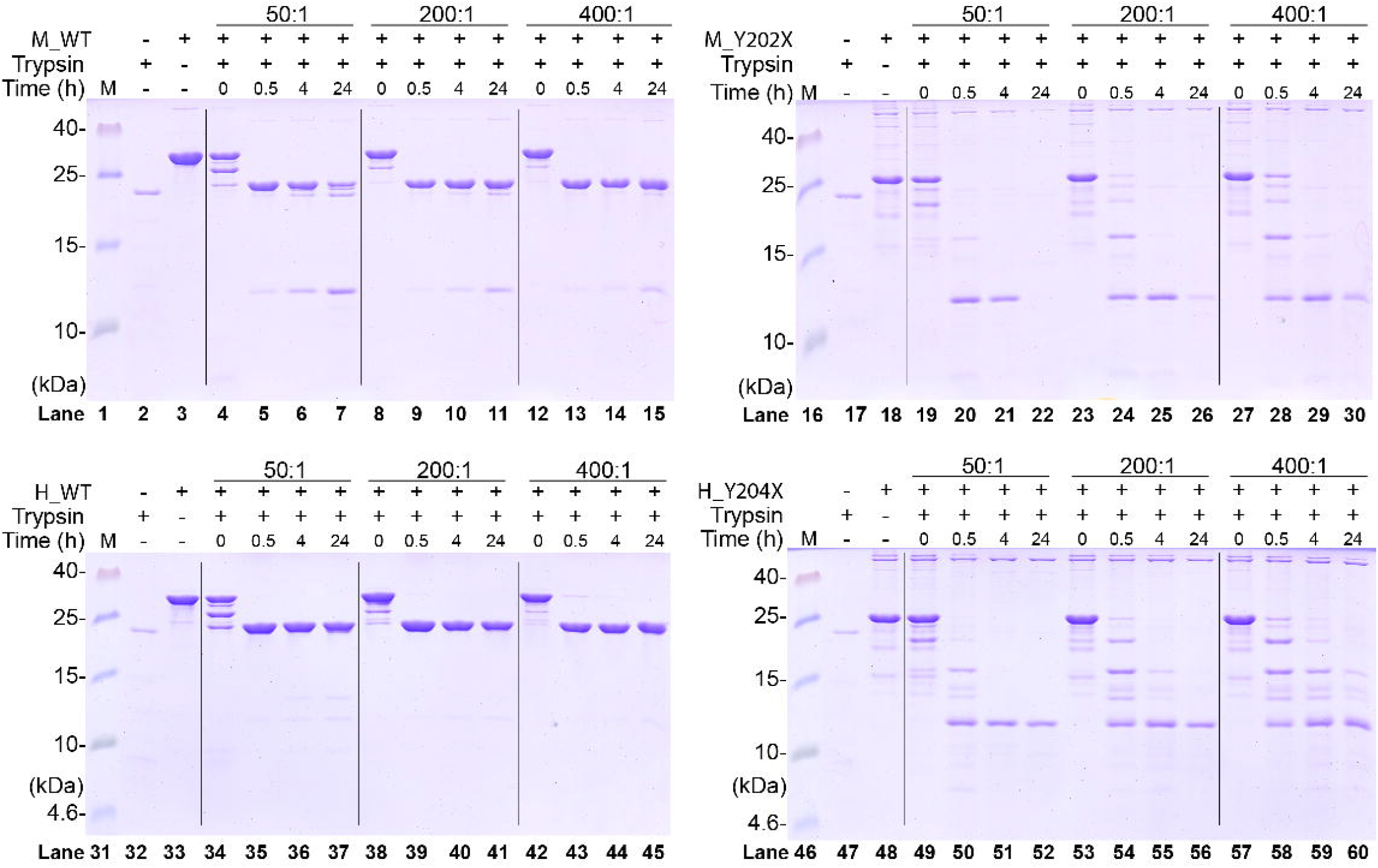
The trypsin proteolysis analyses of the CRYβB1 proteins. The four CRYβB1 proteins (M_WT in lane 1-15, M_Y202X in lane 16-30, H_WT in lane 31-45, H_Y204X in lane 46-60) were treated with protein and trypsin at different ratio (wt. to wt., 50:1, 200:1 and 400:1, divided by black thin line) in a series of time points. The samples at time point 0 h were performed through adding the loading buffer immediately after the trypsin mixing with the CRYβB1 proteins.

## Discussion

According to the public Human Gene Mutation Database and previous reports, 17 *CRY*_Β_*B1* mutations have been associated with the congenital cataract (4,16,17,19-23,25,26). Of them, 8 mutations (including 3 truncation-type mutations) are within CRYβB1 C-terminal region (CTD and C-terminal extension). Previous functional investigation of these CRYβB1 C-terminal region mutations mainly focused on protein solubility, thermal stability, and oligomerization properties, but is not sufficient to elucidate mechanistic details. In this study, we clinically identified a novel cataract-causing CRYβB1 mutation p.Y204X, allowing us to study the mechanism of truncation-type mutations from a different mutation site. Besides DLS and EM data that support alteration in oligomerization propensity caused by the mutation, the M_Y202X NTD crystal structure and trypsin proteolysis analysis together suggest that this mutation disrupts structural integrity of CRYβB1 CTD and could result in complete degradation of the C-terminal region. As suggested by our structural analysis, new NTD-dictating interactions can be established between degraded CRYβB1. We therefore propose a working model for understanding the mechanism of action for CRYβB1 truncation-type mutation (Figure 7). In contrast to WT protein that forms dimer through inter-molecular interactions involving intact NTD and CTD, truncation-type mutants disrupt the structural integrity of CTD either directly or through degradation, thus altering the inter-molecular interaction profiles and may lead to formation of HMW aggregates (Figure 7). As indicated by the M_Y202X NTD structure, complete degradation of the C-terminal region can promote NTD dimer formation. One NTD dimer can well mimic one WT monomer in shape but not in surface properties such as electrostatic features. Once actually formed *in vivo*, the degraded forms of CRYβB1 could modulate inter-molecular interactions between WT CRYβB1 molecules as well as those between CRYβB1 and other crystallins.

**Figure 7.**
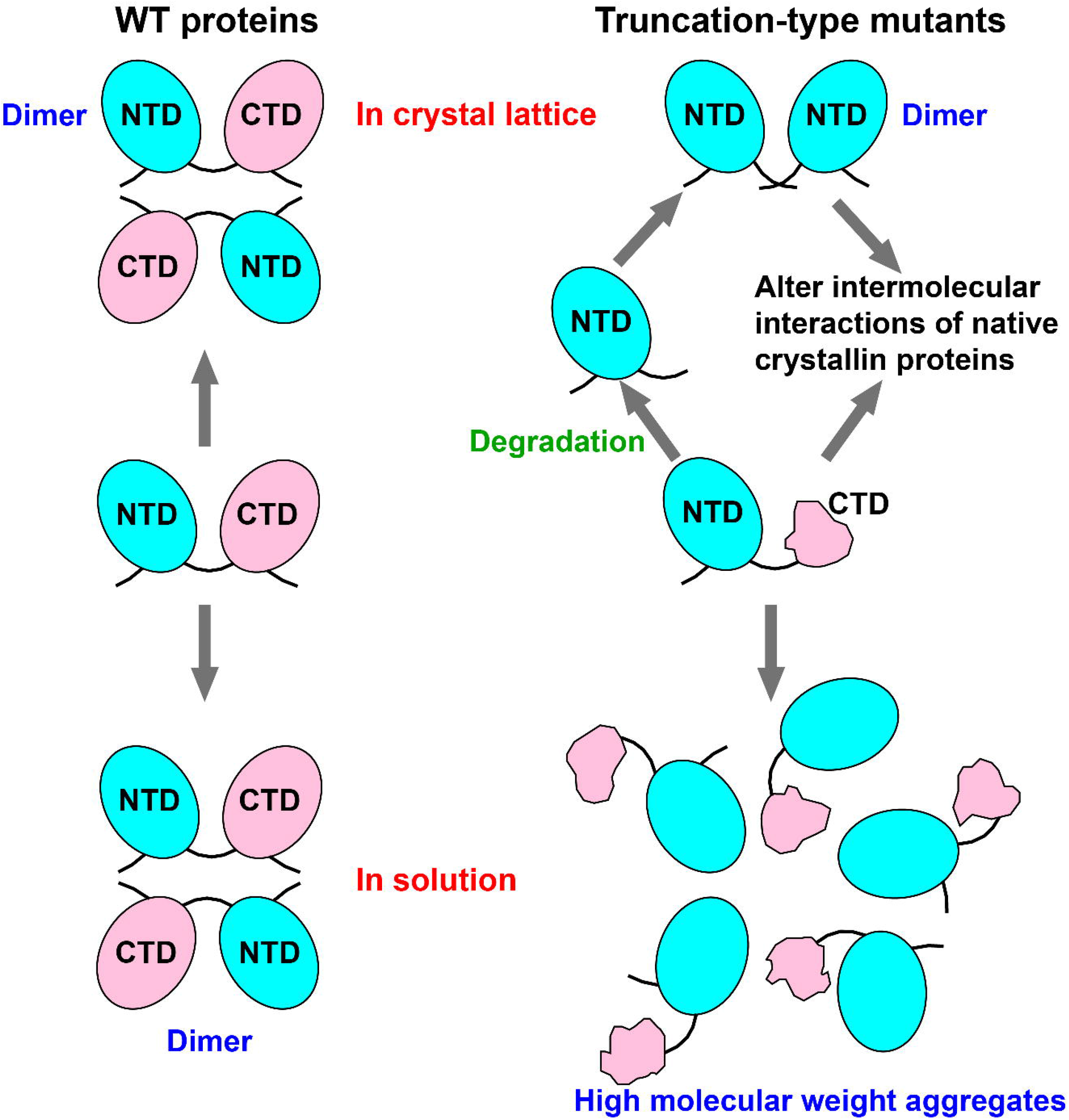
Proposed mechanism of action for *CRYBB1* truncation-type mutations. Left: WT CRYβB1 WT proteins form dimers both in crystal lattice and in solution. Right: CRYβB1 truncation-type mutants (e.g. H_Y204X) form high molecular weight aggregates in solution. These mutants may be susceptible to degradation and become NTD-only variants as captured in CRYβB1 H_Y204X crystal lattice. The NTD-only variant of CRYβB1 can form dimer, and together with the original form of the mutant, may alter inter-molecular interactions of native crystallin proteins. The NTD and CTD of CRYβB1 are colored in cyan and pink, respectively, and the N- and C-terminal extensions are shown as black thin lines. WT proteins: M_WT or H_WT; truncation-type mutants: M_Y202X or H_Y204X.

Two primary models have been proposed for mutation-derived cataract formation (27-29). The first model is featured by mild mutations that do not significantly change protein properties. However, long-term impact from adverse environmental factors may eventually cause protein denaturation and aggregation that affects light scattering and leads to cataract. In the second model, protein structure is disrupted to a great extent by severe mutations that cause congenital cataracts. The truncation-type mutations, including CRYβB1 p.Y204X mutation in this study, belong to the severe mutation category. Here we investigated the molecular properties of congenital cataract-related CRYβB1 p.Y204X mutation in comparison with the WT protein and their counterparts in mouse. Our data, in particular those obtained through crystallography and trypsin proteolytic analysis, suggest that this mutation may change the inter-molecular interactions between ç proteins themselves or between CRYβB1 protein and other crystallins, either through the intact mutant or the degraded NTD dimer that mimics WT monomer (Figure 7). Such mutation-derived alteration of inter-molecular interactions could lead to malfunctioning of lens cells or disruption of crystallin lens microstructures. Our work thus provides an important reference in understanding the structure-function relationship of crystallins and the connection between crystallin mutation and cataract formation.

### Experimental Procedures

#### Ethics

The study protocols were approved by the Human Ethics Committee of the Guangzhou Women and Children’s Medical Center. Written informed consent was obtained from each participant or their legal custodians.

#### Whole exome sequencing

For whole exome sequencing, genomic DNA (gDNA) was extracted from the peripheral blood of all subjects and sonicated into DNA fragments of 150-300 bp. Whole exome sequencing was performed at the Beijing Genomics Institute (Shenzhen, China) as previously described (30). The raw data was collected using Illumina Base Calling software (bcl2fastq). The human genome assembly hg19 (GRCh37) was used as the reference sequence. The Genome Analysis Toolkit (GATK v3.3.0) and ANNOVAR software were employed to detect and annotate the variants, respectively. Then, all variants were filtered following a pipeline: 1) exclusion of variants with a frequency greater than 1% in any of the four databases (1000g_all, esp6500siv2_all, gnomAd_ALL and gnomAD_EAS); 2) exclusion of variants that were not in the coding (exonic) region or splicing region (splicing site ± 10 bp); 3) exclusion of synonymous SNPs that were not predicted by dbscSNV to affect splicing, and 4) retention of variants that were predicted by at least two of four prediction tools (SIFT, Polyphen, MutationTaster, and CADD) to be deleterious and variants that are predicted to affect splicing.

#### Variant verification

Sanger sequencing was performed to validate the pathogenic variant identified by WES. gDNA was used as a template. The following primers were used: 5’-TACCATGCACAGGCAACATGC-3’ (forwards) and 5’-TAGCAGAGTGAGGTGT GGACTC-3’ (reverse). The qualified PCR products were sent to Shanghai Sangon Biotech (Shanghai, China) for sequencing. *CRY*_Β_*B1* reference sequence: NM_001887.4.

#### Plasmid construction, protein expression, and purification

The human CRYβB1 H_WT, human CRYβB1 H_WT, mouse CRYβB1 M_WT, and mouse mutant M_Y202X (codon optimized) genes were cloned into the pET26b vector. The resulting plasmids were transformed into *Escherichia coli* (*E. coli*) strain BL21(DE3) for the expression of the proteins. The cells were cultured in LB medium containing 50 μg/ml for kanamycin at 37 °C until the optical density at 600 nm (OD_600_) reached 0.4-0.6. Then the cultures were induced with isopropyl-β-D-thiogalactopyranoside (IPTG) at a concentration of 0.5 mM at 16 °C overnight. The cells were harvested by centrifugation at 6,740 g for 15 min in an F10S×1000 rotor (Thermo Scientific) and stored at -80 °C. The CRYβB1 proteins purification procedures were modified from previously reported methods with a HiTrap Q/SP HP column (GE Healthcare) used in the second chromatography step and glycerol removed from the gel filtration (GF) buffer (300 mM NaCl, 5 mM Tris pH 7.5) in the third chromatography step (31). The purified protein was supplemented with 5 mM tris-(2-carboxyethyl) phosphine (TCEP) and concentrated to a final concentration of about 3-20 mg/ml, flash frozen with liquid nitrogen, and stored at -80 °C as 10-20 μl aliquots for single use. The extinction coefficient of CRYβB1 proteins were calculated based on the amino acid sequences using the ExPASy ProtParam program (http://www.expasy.ch/tools/protparam.html).

#### Crystallization, data collection, and structure determination

The crystallization screening of CRYβB1 proteins was performed at 289 K by sitting drop vapor diffusion. The M_Y202X plate-like crystal yielding the final X-ray diffraction dataset was obtained by mixing 0.3 μl of 10 mg/ml M_Y202X, 0.3 μl of a reservoir solution of 0.1 M sodium malonate (pH 6.0) and 12% (w/v) Polyethylene glycol 3350. Crystals were soaked in the precipitant solution supplemented with 26.7% (vol./vol.) glycerol as a cryo-protectant before being flash-cooled and stored in liquid nitrogen. The X-ray diffraction data was collected at the Shanghai Synchrotron Radiation Facility (SSRF) beamline BL17U1 (wavelength = 0.9792 Å, temperature = 100 K) (32). Reflections were integrated, merged, and scaled using HKL2000 (Table 1) (33). The initial structure was obtained using the molecular replacement program PHASER (34) with a truncated human CRYβB1 crystal structure (PDB entry 1OKI) (1) as a search model with the C-terminal domain removed. Manual model rebuilding and refinement was completed by Coot (35) and Phenix, respectively (36). Non-crystallographic (NCS) symmetry was applied to both chains in the asymmetric unit in initial rounds of refinement and was released in later rounds. The Ramachandran statistics are 89.3%, 10.7%, 0.0%, and 0.0% for favored, allowed, generously allowed, and disfavored regions, respectively. The 3,500 K composite simulated-annealing (SA) omit 2F_o_-F_c_ electron density maps were generated by CNS (37). All structure superimpositions were performed using the maximum likelihood structure superpositioning program THESEUS (38). The pdb entry used for superpositioning analysis with CRY βB1 Y202X is 1OKI (CRY βB1 WT) (1).

#### Measurement of hydrodynamic size

In order to compare the aggregation tendency of the CRYβB1 proteins in solution, the H_WT, H_Y204X, M_WT and M_Y202X protein samples in PBS were centrifuged at 15,871 *g* in a 5424/5424R rotor (Eppendorf) for 20 min at 25 °C. The supernatants of the samples were used for the measurement of the hydrodynamic size of the protein particles. Measurements were performed on a Zetasizer Nano ZS90 (Malvern Panalytical Ltd, UK) and the parameters were automatically optimized. Each sample was measured three times.

#### Negative-staining analysis

For negative-staining assays, the CRYβB1 proteins were diluted in the GF buffer to 0.02 mg/mL. 10 μl of protein sample was loaded onto a glow-discharged carbon-coated copper grid and stained with 3% Uranyl Acetate (UA). The prepared grids were examined using an FEI Tecnai 20 TEM (operated at 200 kV) equipped with a Gatan UltraScan 894 CCD camera. The TEM images were trimed using Photoshop software without any brightness/contrast adjustment.

#### Trypsin proteolysis assays

A 20 μl reaction mixture containing protein samples and trypsin at 50:1, 200:1 and 400:1 (w/w) in 25 mM Tris-HCl (pH 8.0) was incubated at 25 °C for 0, 0.5, 4 and 24 h before being boiled with an equal volume of Tris-Tricine-SDS-PAGE loading buffer (40 mM Tris-HCl, pH 8.0, 40% (v/v) glycerol, 20 mM DTT, 4% (w/v) SDS, 0.02% (w/v/) bromophenol blue). The samples were analyzed by Tris-Tricine-SDS-PAGE.

### Data Availability

All data acquired are available upon request.

## Acknowledgements

This study was supported by the National Natural Science Foundation of China (82101102 to X.J.), the Key Biosafety Science and Technology Program of Hubei Jiangxia Laboratory (JXBS001 to P.G.), and the Natural Science Foundation of Guangdong Province, China (2022A1515012506 to M.Z.). We thank synchrotrons SSRF (beamline BL17U1/BL10U2 Shanghai, China) for access to beamline, Dr. Jiqin Wu, Xiang Fang, Qiaojie Liu, Tianying Nong, and Caixia Xian for laboratory assistance.

## Contributions

Y.-P.T. and P.G. conceived the study; X.L., M.Z., L.S., and P.W. obtained clinical and genetics data; X.J. and B.-Y.Z. performed the crystallographic and biochemical experiments; Y.X., D.-M.X., and Y.-P.T. analyzed the clinical and genetics data; X.J., B.-Y.Z., and P.G. analyzed the crystallographic and biochemical data; X.J., M.Z., Y.-P.T. and P.G. wrote the paper; all authors approved the final version of the manuscript.

## Accession number

The atomic coordinates and structure factor files have been deposited in the Protein Data Bank under accession code 8H0R.

