## Supplemental file 1 for "A cataract-causing Y204X mutation of CRYßB1 promotes C-terminal degradation and higher-order oligomerization"

### Full wwPDB X-ray Structure Validation Report (i)

Oct 7, 2022 – 10:54 AM JST

| | | |
|---|---|---|
| PDB ID | : | 8H0R |
| Title | : | Crystal Structure of a novel cataract-causing human crystallin CRYBB1 truncated mutant Y204X |
| Deposited on | : | 2022-09-30 |
| Resolution | : | 1.20 Å(reported) |

**This wwPDB validation report is for manuscript review**

This is a Full wwPDB X-ray Structure Validation Report.

This report is produced by the wwPDB biocuration pipeline after annotation of the structure.

We welcome your comments at **
A user guide is available at
https://www.wwpdb.org/validation/2017/XrayValidationReportHelp
with specific help available everywhere you see the (i) symbol.

The types of validation reports are described at
http://www.wwpdb.org/validation/2017/FAQs#types.

For Manuscript Review

The following versions of software and data (see references (i)) were used in the production of this report:

| | | |
|---|---|---|
| MolProbity | : | 4.02b-467 |
| Xtriage (Phenix) | : | 1.13 |
| EDS | : | 2.31.2 |
| Percentile statistics | : | 20191225.v01 (using entries in the PDB archive December 25th 2019) |
| Refmac | : | 5.8.0158 |
| CCP4 | : | 7.0.044 (Gargrove) |
| Ideal geometry (proteins) | : | Engh & Huber (2001) |
| Ideal geometry (DNA, RNA) | : | Parkinson et al. (1996) |
| Validation Pipeline (wwPDB-VP) | : | 2.31.2 |

### 1 Overall quality at a glance (i)

The following experimental techniques were used to determine the structure:
*X-RAY DIFFRACTION*

The reported resolution of this entry is 1.20 Å.

Percentile scores (ranging between 0-100) for global validation metrics of the entry are shown in the following graphic. The table shows the number of entries on which the scores are based.

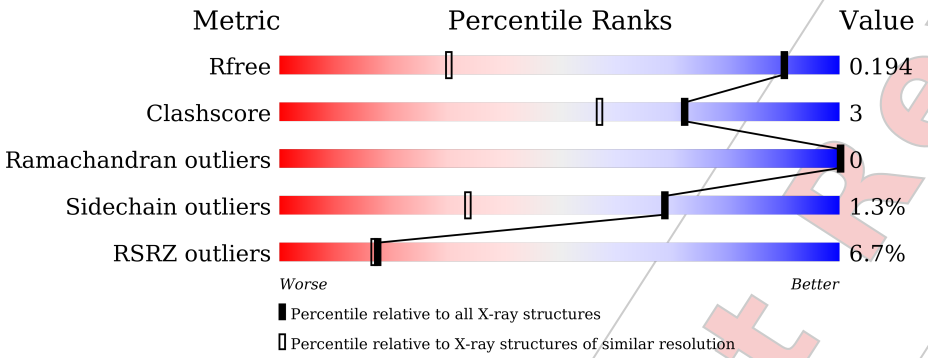


| Metric | Whole archive (#Entries) | Similar resolution (#Entries, resolution range(Å)) |
|---|---|---|
| $R_{free}$ | 130704 | 1223 (1.22-1.18) |
| Clashscore | 141614 | 1286 (1.22-1.18) |
| Ramachandran outliers | 138981 | 1240 (1.22-1.18) |
| Sidechain outliers | 138945 | 1239 (1.22-1.18) |
| RSRZ outliers | 127900 | 1200 (1.22-1.18) |

The table below summarises the geometric issues observed across the polymeric chains and their fit to the electron density. The red, orange, yellow and green segments of the lower bar indicate the fraction of residues that contain outliers for >=3, 2, 1 and 0 types of geometric quality criteria respectively. A grey segment represents the fraction of residues that are not modelled. The numeric value for each fraction is indicated below the corresponding segment, with a dot representing fractions <=5% The upper red bar (where present) indicates the fraction of residues that have poor fit to the electron density. The numeric value is given above the bar.

| Mol | Chain | Length | Quality of chain |
|---|---|---|---|
| 1 | A | 211 | 3%<br>39% • 57% |
| 1 | B | 211 | 2%<br>39% • 59% |

For Manuscript Review

### 2 Entry composition (i)

There are 2 unique types of molecules in this entry. The entry contains 1694 atoms, of which 0 are hydrogens and 0 are deuteriums.

In the tables below, the ZeroOcc column contains the number of atoms modelled with zero occupancy, the AltConf column contains the number of residues with at least one atom in alternate conformation and the Trace column contains the number of residues modelled with at most 2 atoms.

- Molecule 1 is a protein called Beta-crystallin B1.

| Mol | Chain | Residues | Atoms | | | | | ZeroOcc | AltConf | Trace |
|---|---|---|---|---|---|---|---|---|---|---|
| 1 | A | 91 | Total 766 | C 489 | N 131 | O 141 | S 5 | 0 | 4 | 0 |
| 1 | B | 87 | Total 741 | C 476 | N 130 | O 132 | S 3 | 0 | 4 | 0 |

There are 18 discrepancies between the modelled and reference sequences:

| Chain | Residue | Modelled | Actual | Comment | Reference |
|---|---|---|---|---|---|
| A | -7 | MET | - | initiating methionine | UNP Q9WVJ5 |
| A | -6 | HIS | - | expression tag | UNP Q9WVJ5 |
| A | -5 | HIS | - | expression tag | UNP Q9WVJ5 |
| A | -4 | HIS | - | expression tag | UNP Q9WVJ5 |
| A | -3 | HIS | - | expression tag | UNP Q9WVJ5 |
| A | -2 | HIS | - | expression tag | UNP Q9WVJ5 |
| A | -1 | HIS | - | expression tag | UNP Q9WVJ5 |
| A | 0 | GLY | - | expression tag | UNP Q9WVJ5 |
| A | 1 | SER | - | expression tag | UNP Q9WVJ5 |
| B | -7 | MET | - | initiating methionine | UNP Q9WVJ5 |
| B | -6 | HIS | - | expression tag | UNP Q9WVJ5 |
| B | -5 | HIS | - | expression tag | UNP Q9WVJ5 |
| B | -4 | HIS | - | expression tag | UNP Q9WVJ5 |
| B | -3 | HIS | - | expression tag | UNP Q9WVJ5 |
| B | -2 | HIS | - | expression tag | UNP Q9WVJ5 |
| B | -1 | HIS | - | expression tag | UNP Q9WVJ5 |
| B | 0 | GLY | - | expression tag | UNP Q9WVJ5 |
| B | 1 | SER | - | expression tag | UNP Q9WVJ5 |

- Molecule 2 is water.

| Mol | Chain | Residues | Atoms | | ZeroOcc | AltConf |
|---|---|---|---|---|---|---|
| 2 | A | 102 | Total 102 | O 102 | 0 | 0 |

*Continued on next page...*

For Manuscript Review

*Continued from previous page...*

| Mol | Chain | Residues | Atoms | ZeroOcc | AltConf |
|---|---|---|---|---|---|
| 2 | B | 85 | Total O<br>85 85 | 0 | 0 |

For Manuscript Review

### 3 Residue-property plots (i)

These plots are drawn for all protein, RNA, DNA and oligosaccharide chains in the entry. The first graphic for a chain summarises the proportions of the various outlier classes displayed in the second graphic. The second graphic shows the sequence view annotated by issues in geometry and electron density. Residues are color-coded according to the number of geometric quality criteria for which they contain at least one outlier: green = 0, yellow = 1, orange = 2 and red = 3 or more. A red dot above a residue indicates a poor fit to the electron density (RSRZ > 2). Stretches of 2 or more consecutive residues without any outlier are shown as a green connector. Residues present in the sample, but not in the model, are shown in grey.

• Molecule 1: Beta-crystallin B1

Chain A:

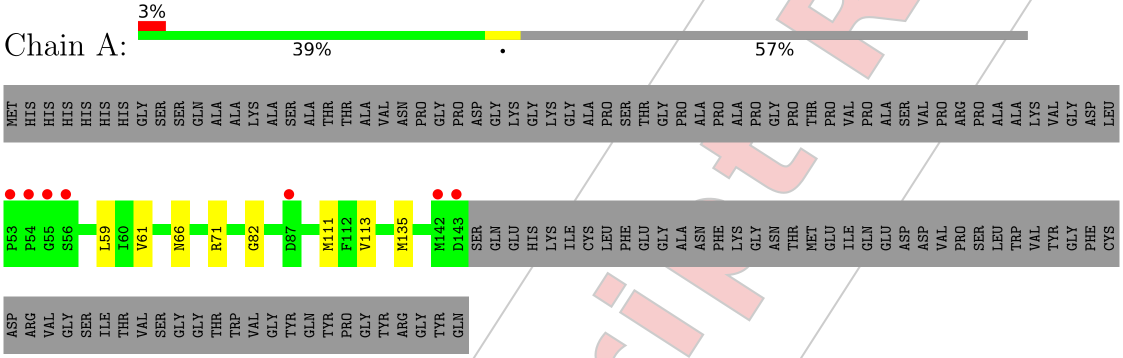


• Molecule 1: Beta-crystallin B1

Chain B:

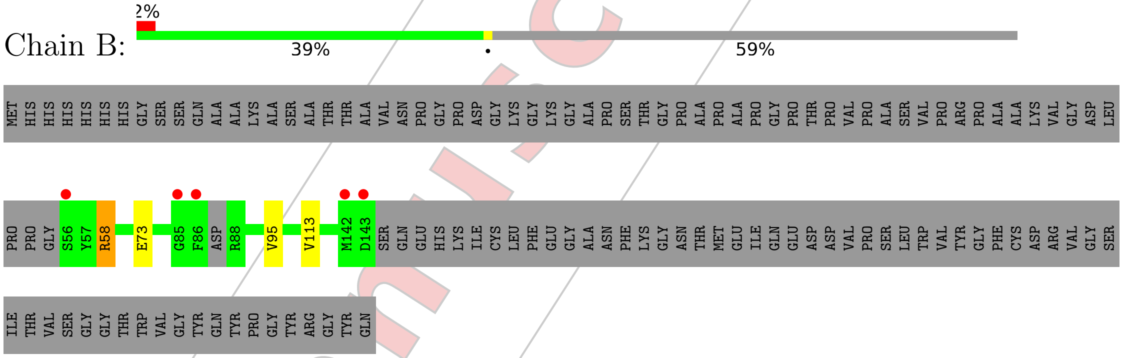


For Manuscript Review

### 4 Data and refinement statistics (i)

| Property | Value | Source |
|---|---|---|
| Space group | P 21 21 21 | Depositor |
| Cell constants<br>a, b, c, $\alpha$, $\beta$, $\gamma$ | 39.92Å 48.44Å 113.32Å<br>90.00° 90.00° 90.00° | Depositor |
| Resolution (Å) | 23.10 – 1.20<br>36.82 – 1.20 | Depositor<br>EDS |
| % Data completeness<br>(in resolution range) | 98.0 (23.10-1.20)<br>98.0 (36.82-1.20) | Depositor<br>EDS |
| $R_{merge}$ | 0.06 | Depositor |
| $R_{sym}$ | (Not available) | Depositor |
| $< I/\sigma(I) >$ [1] | 2.21 (at 1.20Å) | Xtriage |
| Refinement program | PHENIX 1.12 | Depositor |
| R, $R_{free}$ | 0.185 , 0.194<br>0.185 , 0.194 | Depositor<br>DCC |
| $R_{free}$ test set | 3398 reflections (4.99%) | wwPDB-VP |
| Wilson B-factor (Å$^2$) | 14.4 | Xtriage |
| Anisotropy | 0.153 | Xtriage |
| Bulk solvent $k_{sol}$(e/Å$^3$), $B_{sol}$(Å$^2$) | 0.39 , 40.4 | EDS |
| L-test for twinning[2] | $< \|L\| > = 0.49$, $< L^2 > = 0.32$ | Xtriage |
| Estimated twinning fraction | No twinning to report. | Xtriage |
| $F_o$,$F_c$ correlation | 0.97 | EDS |
| Total number of atoms | 1694 | wwPDB-VP |
| Average B, all atoms (Å$^2$) | 19.0 | wwPDB-VP |

Xtriage's analysis on translational NCS is as follows: *The largest off-origin peak in the Patterson function is 8.50% of the height of the origin peak. No significant pseudotranslation is detected.*

[1] Intensities estimated from amplitudes.

[2] Theoretical values of $< |L| >$, $< L^2 >$ for acentric reflections are 0.5, 0.333 respectively for untwinned datasets, and 0.375, 0.2 for perfectly twinned datasets.

For Manuscript Review

### 5 Model quality (i)

#### 5.1 Standard geometry (i)

The Z score for a bond length (or angle) is the number of standard deviations the observed value is removed from the expected value. A bond length (or angle) with $|Z| > 5$ is considered an outlier worth inspection. RMSZ is the root-mean-square of all Z scores of the bond lengths (or angles).

| Mol | Chain | Bond lengths | | Bond angles | |
|---|---|---|---|---|---|
| | | RMSZ | #$\|Z\|$ >5 | RMSZ | #$\|Z\|$ >5 |
| 1 | A | 0.37 | 0/796 | 0.62 | 0/1072 |
| 1 | B | 0.40 | 0/771 | 0.62 | 0/1037 |
| All | All | 0.39 | 0/1567 | 0.62 | 0/2109 |

There are no bond length outliers.

There are no bond angle outliers.

There are no chirality outliers.

There are no planarity outliers.

#### 5.2 Too-close contacts (i)

In the following table, the Non-H and H(model) columns list the number of non-hydrogen atoms and hydrogen atoms in the chain respectively. The H(added) column lists the number of hydrogen atoms added and optimized by MolProbity. The Clashes column lists the number of clashes within the asymmetric unit, whereas Symm-Clashes lists symmetry-related clashes.

| Mol | Chain | Non-H | H(model) | H(added) | Clashes | Symm-Clashes |
|---|---|---|---|---|---|---|
| 1 | A | 766 | 0 | 726 | 7 | 0 |
| 1 | B | 741 | 0 | 716 | 6 | 0 |
| 2 | A | 102 | 0 | 0 | 1 | 0 |
| 2 | B | 85 | 0 | 0 | 0 | 0 |
| All | All | 1694 | 0 | 1442 | 10 | 0 |

The all-atom clashscore is defined as the number of clashes found per 1000 atoms (including hydrogen atoms). The all-atom clashscore for this structure is 3.

All (10) close contacts within the same asymmetric unit are listed below, sorted by their clash magnitude.

| Atom-1 | Atom-2 | Interatomic distance (Å) | Clash overlap (Å) |
|---|---|---|---|
| 1:B:58:ARG:NH1 | 1:B:73:GLU:OE1 | 2.33 | 0.61 |

*Continued on next page...*

For Manuscript Review

*Continued from previous page...*

| Atom-1 | Atom-2 | Interatomic distance (Å) | Clash overlap (Å) |
|---|---|---|---|
| 1:B:58:ARG:HE | 1:B:73:GLU:CD | 2.03 | 0.61 |
| 1:B:58:ARG:HB3 | 1:B:95:VAL:HB | 1.83 | 0.60 |
| 1:A:113:VAL:HG22 | 1:B:113[A]:VAL:HG22 | 1.92 | 0.52 |
| 1:A:113:VAL:CG2 | 1:B:113[A]:VAL:HG22 | 2.43 | 0.48 |
| 1:A:66:ASN:HB3 | 2:A:387:HOH:O | 2.15 | 0.47 |
| 1:A:59:LEU:HD21 | 1:A:61:VAL:HG13 | 1.99 | 0.44 |
| 1:A:59:LEU:CD2 | 1:A:61:VAL:HG13 | 2.49 | 0.43 |
| 1:A:82:GLY:H | 1:A:135[B]:MET:HE1 | 1.85 | 0.42 |
| 1:A:111:MET:SD | 1:B:113[B]:VAL:HG12 | 2.61 | 0.41 |

There are no symmetry-related clashes.

#### 5.3 Torsion angles (i)

##### 5.3.1 Protein backbone (i)

In the following table, the Percentiles column shows the percent Ramachandran outliers of the chain as a percentile score with respect to all X-ray entries followed by that with respect to entries of similar resolution.

The Analysed column shows the number of residues for which the backbone conformation was analysed, and the total number of residues.

| Mol | Chain | Analysed | Favoured | Allowed | Outliers | Percentiles | |
|---|---|---|---|---|---|---|---|
| 1 | A | 93/211 (44%) | 92 (99%) | 1 (1%) | 0 | 100 | 100 |
| 1 | B | 87/211 (41%) | 85 (98%) | 2 (2%) | 0 | 100 | 100 |
| All | All | 180/422 (43%) | 177 (98%) | 3 (2%) | 0 | 100 | 100 |

There are no Ramachandran outliers to report.

##### 5.3.2 Protein sidechains (i)

In the following table, the Percentiles column shows the percent sidechain outliers of the chain as a percentile score with respect to all X-ray entries followed by that with respect to entries of similar resolution.

The Analysed column shows the number of residues for which the sidechain conformation was analysed, and the total number of residues.

For Manuscript Review

| Mol | Chain | Analysed | Rotameric | Outliers | Percentiles | |
|---|---|---|---|---|---|---|
| 1 | A | 83/172 (48%) | 82 (99%) | 1 (1%) | 71 | 37 |
| 1 | B | 80/172 (46%) | 79 (99%) | 1 (1%) | 69 | 33 |
| All | All | 163/344 (47%) | 161 (99%) | 2 (1%) | 69 | 37 |

All (2) residues with a non-rotameric sidechain are listed below:

| Mol | Chain | Res | Type |
|---|---|---|---|
| 1 | A | 71 | ARG |
| 1 | B | 58 | ARG |

Sometimes sidechains can be flipped to improve hydrogen bonding and reduce clashes. There are no such sidechains identified.

##### 5.3.3 RNA (i)

There are no RNA molecules in this entry.

#### 5.4 Non-standard residues in protein, DNA, RNA chains (i)

There are no non-standard protein/DNA/RNA residues in this entry.

#### 5.5 Carbohydrates (i)

There are no monosaccharides in this entry.

#### 5.6 Ligand geometry (i)

There are no ligands in this entry.

#### 5.7 Other polymers (i)

There are no such residues in this entry.

#### 5.8 Polymer linkage issues (i)

There are no chain breaks in this entry.

For Manuscript Review

### 6 Fit of model and data (i)

#### 6.1 Protein, DNA and RNA chains (i)

In the following table, the column labelled '#RSRZ> 2' contains the number (and percentage) of RSRZ outliers, followed by percent RSRZ outliers for the chain as percentile scores relative to all X-ray entries and entries of similar resolution. The OWAB column contains the minimum, median, $95^{th}$ percentile and maximum values of the occupancy-weighted average B-factor per residue. The column labelled 'Q< 0.9' lists the number of (and percentage) of residues with an average occupancy less than 0.9.

| Mol | Chain | Analysed | <RSRZ> | #RSRZ>2 | | | OWAB(Å$^2$) | Q<0.9 |
|---|---|---|---|---|---|---|---|---|
| 1 | A | 91/211 (43%) | 0.56 | 7 (7%) | 13 | 12 | 11, 15, 31, 41 | 0 |
| 1 | B | 87/211 (41%) | 0.50 | 5 (5%) | 23 | 22 | 12, 17, 35, 45 | 0 |
| All | All | 178/422 (42%) | 0.53 | 12 (6%) | 17 | 17 | 11, 16, 36, 45 | 0 |

All (12) RSRZ outliers are listed below:

| Mol | Chain | Res | Type | RSRZ |
|---|---|---|---|---|
| 1 | B | 85 | GLY | 7.0 |
| 1 | A | 143 | ASP | 6.2 |
| 1 | A | 53 | PRO | 5.8 |
| 1 | B | 86 | PHE | 4.9 |
| 1 | A | 54 | PRO | 4.4 |
| 1 | B | 143 | ASP | 4.2 |
| 1 | B | 56 | SER | 4.0 |
| 1 | A | 55 | GLY | 3.8 |
| 1 | A | 142 | MET | 3.3 |
| 1 | B | 142 | MET | 2.7 |
| 1 | A | 87 | ASP | 2.7 |
| 1 | A | 56 | SER | 2.4 |

#### 6.2 Non-standard residues in protein, DNA, RNA chains (i)

There are no non-standard protein/DNA/RNA residues in this entry.

#### 6.3 Carbohydrates (i)

There are no monosaccharides in this entry.

For Manuscript Review

#### 6.4 Ligands (i)

There are no ligands in this entry.

#### 6.5 Other polymers (i)

There are no such residues in this entry.

For Manuscript Review
